# *Pseudolaguvia moinensis*, a new benthic miniature Asian catfish (Teleostei: Sisoridae) from Arunachal Pradesh, north-eastern India

**DOI:** 10.1101/2021.09.27.461973

**Authors:** Lakpa Tamang, Prasanta Nanda, Debangshu Narayan Das

## Abstract

A new species of miniature sisorid catfish is described, from the upper Brahmaputra river drainage in northeastern India. The new species is distinguished from all its congeners except *P. ferruginea* and *P. focusa* by having an elongate light brown to cream marks either side on ventro-lateral margin just above anus. Further, distinguished from its congeners by the following combination of characters: presence of a W-shaped dark brown band on caudal fin, thoracic adhesive apparatus extending closer to pelvic-fin base, smooth anterior margin of dorsal spine, a narrow V-shaped light brown to cream bands on side of the body, dorsal-spine length (10.7–14.7% SL), dorsal-fin base length (10.7–14.2% SL), pectoral-fin spine length (14.0–21.1% SL), pelvic-fin length (13.5–16.6% SL), caudal peduncle length (14.4–18.2% SL) and depth (7.8–9.9% SL), total vertebrae (30–31), caudal fin with a complete medial hyaline bands towards its anterior end, reaching outer margin of each lobe. Other combination of characters differentiating the new species from its congeners are provided in the respective diagnoses.

## Introduction

Members of the south Asian sisorid catfish genus *Pseudolaguvia* are small sized benthic fishes, recorded not more than 47 mm SL, and occurs throughout hill streams and larger rivers along the sub-Himalayan foothills, spanning from the Ganges River drainage, Nepal in the west, Ganges-Brahmaputra River basin in the north and northeast India, the Bharatapuzha River, Western Ghats, Kerala, and the Kumaradhara River, Karnataka, in the south, the Brahmaputra drainages in Bangladesh and the Ayeyarwaddy and Sittang drainages in Myanmar in the east (Ng 2006a; Ng & Lalramliana 2010a; Ng & Lalramliana 2010b; Radhakrishnan *et al*., 2011; Britz *et al*., 2013; Ng & Conway, 2013; Rayamajhi *et al*., 2016).

Currently, twenty three species of *Pseudolaguvia* are recognized (Ng *et al*., 2016; Bhakat, 2019): *P. ribeiroi* (Hora, 1921), *P. shawi* (Hora, 1921), *P. foveolata* Ng, 2005, *P. ferula* Ng, 2006, *P. ferruginea* Ng, 2009, *P. viriosa* Ng & Tamang, 2012, *P. magna* Tamang & Sinha, 2014, *P. jiyaensis* Tamang & Sinha, 2014 and *P. muricata* Ng, 2005, *P. kapuri* (Tilak & Hussain, 1973), *P. inornata* Ng, 2005, *P. flavida* Ng, 2009, *P. virgulata* Ng & Lalramliana, 2010, *P. assula* Ng & Conway, 2013, *P. nepalensis* Rayamajhi *et al*., 2016, *P. flavipinna* Bhakat, 2019, *P. nubila* Ng *et al*., 2013, and *P. fucosa* Ng *et al*., 2016, *P. spicula* Ng & Lalramliana, 2010, *P. austrina* Radhakrishnan, *et al*. 2011, *P. lapillicola* Britz *et al*., 2013, *P. tenebricosa* Britz & Ferraris, 2013 and *P. tuberculata* (Prashad & Mukerji, 1929).

While conducting icththyological surveys in relation to DBT New Delhi sponsored project at Moin drainage about 10 km away in the western corner of the capital city, Itanagar, Arunachal Pradesh, resulted in the collection of thirteen specimens (27.0–31.7 mm SL) of *Pseudolaguvia* along with other targeted species required in the project. After examination and comparison of this material with known congeners revealed it to represent an undescribed species, described in this paper as *Pseudolaguvia moinensis*, new species, and is the fourth report of *Pseudolaguvia* from the foothills of the state Arunachal Pradesh.

## Material and methods

One course of the bifurcated drainage was blocked using 5 m long plastic sheet, sand and soil, diverting water to another course. Fishes were collected by hand as well as with mosquito net along the blockade course. A 2 m diameter nylon cast net with an eye-size of 7×7 mm was also additionally used for sampling fish nearby. Specimens were preserved in 5% formalin at the beginning and later transferred to 75% alcohol. Measurements were made point to point with digital calipers and data recorded to the nearest 0.1 mm. All measurements and counts were made on the left side of specimens whenever possible, following Ng & Kottelat (2013). Subunits of the head are presented as a percentage of head length (HL), and head length and measurements of body parts as a percentage of standard length (SL). For vertebral counts, flesh was removed from the right side (4 specimens: 27.2–28.9 mm SL) using a scalpel and needle, and counted from anterior most weberian vertebrae to last complete vertebrae at caudal-fin base. Fin rays, pectoral-fin spine serrae, vertebrae and number of pores along the lateral line were counted under a stereozoom transmitted light microscope (Magnus: MS-24). The number of individuals exhibiting a given meristic value is indicated in parentheses. Morphometric data are expressed as percentage of standard length for part of the body and head length for parts of the head. Values for the holotype are indicated by an asterisk in the text. Abbreviations used: DBT, Department of Biotechnology, RGUMF, Rajiv Gandhi University Museum of Fishes, Rono Hills, Doimukh, ZSI, Zoological Survey of India, APRC, Arunachal Pradesh Regional Centre, and DNGCMF, Dera Natung Govt College Museum of Fishes, Itanagar, Arunachal Pradesh, India.

### *Pseudolaguvia moinensis*. sp. nov

(Fig. 1)

**FIGURE 1.**
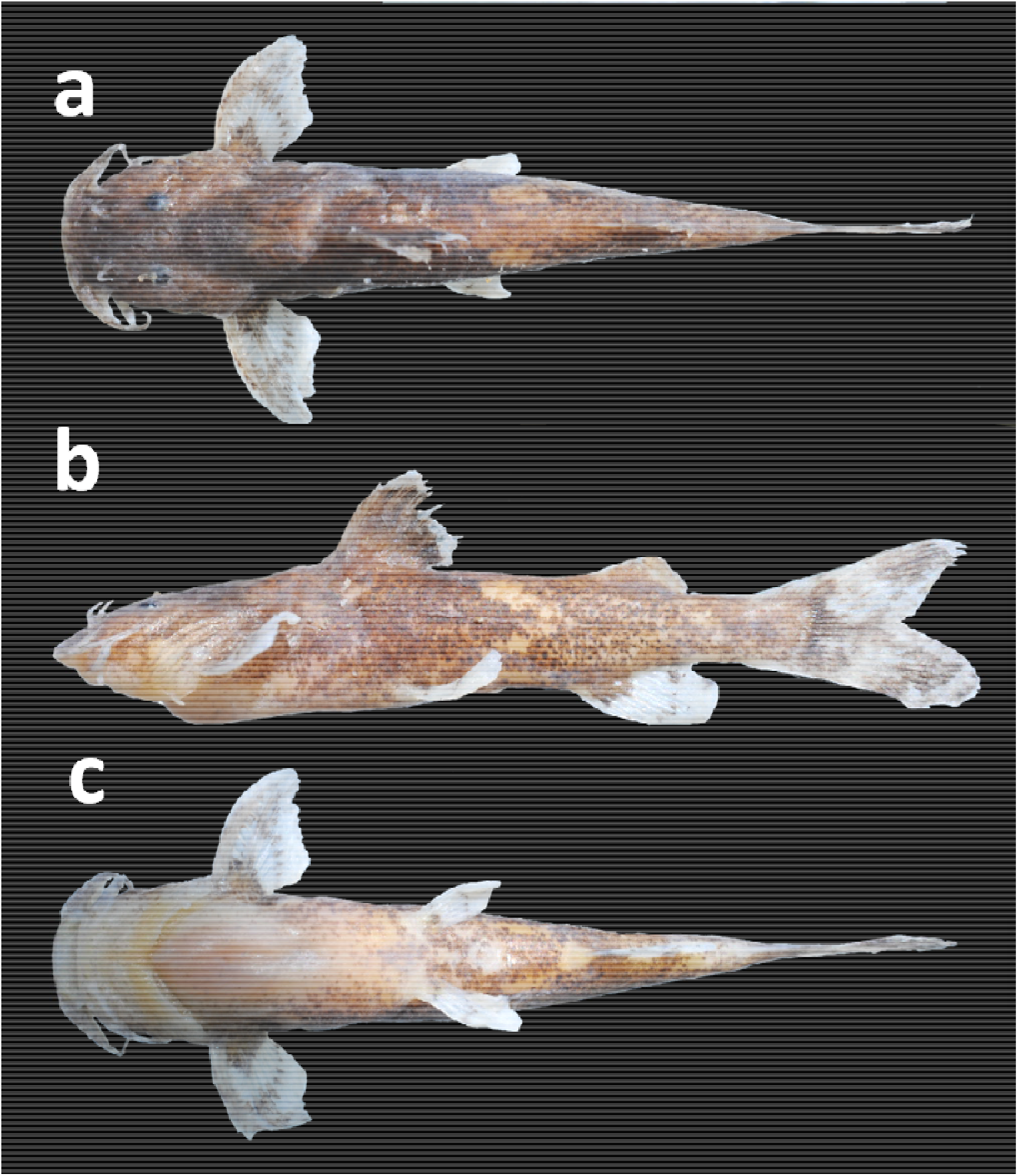
*Pseudolaguvia moinensis*, RGUMF 0553, holotype, 31.4 mm SL; (a) dorsal, (b) lateral and (c) ventral views.

**FIGURE2.**
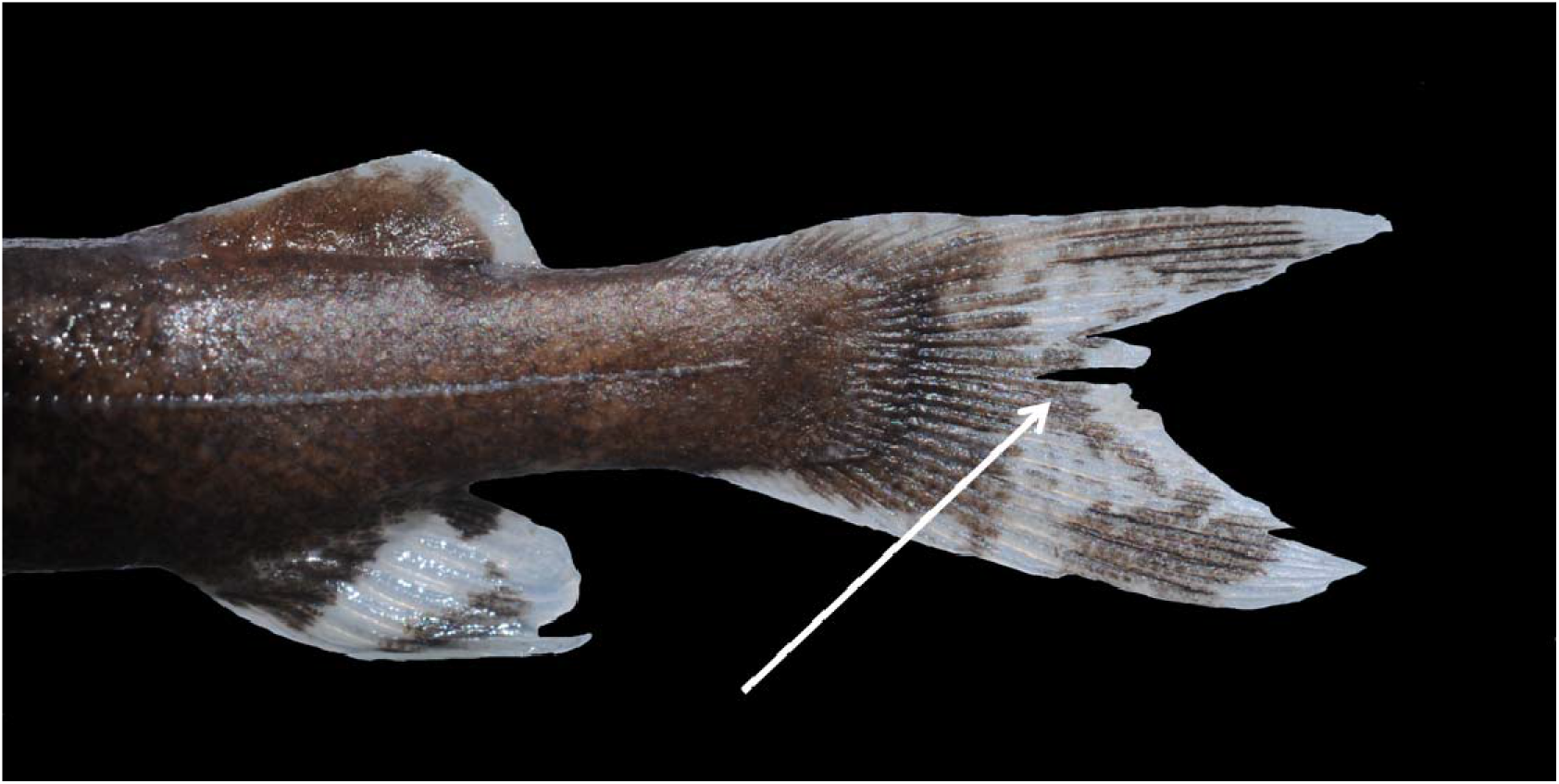
*Pseudolaguviamoinensis*, DNGCMF 06, 28.3 mm SL, caudal fin showing dark brown ‘W’ band.

### Type material

Holotype: ZSI/APRC/F 1897, 31.4 mm SL; India: Arunachal Pradesh, Papum Pare district, Moin drainage, a tributary of Poma River (Brahmaputra River basin), 27° 03□59□N 93°32□17□E; altitude 315 m asl; L. Tamang & party, 25 September 2020.

#### Paratypes

RGUMF 0554, 03, 27.0–30.0 mm SL; RGUMF 0556, 06 (04 specimens used for vertebrae), 27.2–29.9 mm SL; DNGCMF 06, 03, 28.3–31.7 mm SL; 28-30 September, 2020, other data as for holotype.

#### Diagnosis

The new species is diagnosed from all congeners except *P. ferruginea* and *P. focusa* by having an elongate light brown to cream marks either side on ventro-lateral margin just above anus. Further can be diagnosed by the following combination of characters: presence of a W-shaped dark brown band on caudal fin, thoracic adhesive apparatus extending closer to pelvic-fin base, smooth anterior margin of dorsal spine, a narrow V-shaped light brown to cream bands along the flank of the body, dorsal-spine length (10.7–14.7% SL), dorsal-fin base length (10.7–14.2% SL), pectoral-fin spine length (14.0–21.1% SL), pelvic-fin length (13.5–16.6% SL), caudal peduncle length (14.4–18.2% SL) and depth (7.8–9.9% SL), total vertebrae (30–31), caudal fin with a complete medial hyaline bands towards its anterior end, touching outer margin of each lobe. Other few combination of characters differentiating the new species from its congeners are provided in the respective diagnoses.

#### Description

General body shape as in Figure 1; biometric and meristic data are provided in Table 1 and Table 2 respectively. Body moderately elongate, deepest at dorsal-fin origin, deeper than wide, moderately compressed anteriorly, degree of compression increasing from insertion of dorsal fin to caudal-fin base. Dorsal profile overall evenly rising to dorsal-fin origin, and then gently decreasing to caudal-fin base. Ventral profile almost horizontal upto to pelvic-fin origin, and then very gently ascending to caudal-fin base.

**Table 1.** Biometric data of *Pseudolaguvia moinensis* (n=13). Range include data of holotype.

| Characters | Holotype | Range | Means±SD |
| --- | --- | --- | --- |
| Standard length (mm) | 31.4 | 27.0–31.7 | 27.1±1.5 |
| %SL |  |  |  |
| Predorsal length | 38.1 | 38.1–42.9 | 37.9±1.4 |
| Preanal length | 67.0 | 65.8–72.1 | 64.9±1.9 |
| Prepelvic length | 47.9 | 47.1–52.8 | 46.8±1.7 |
| Prepectoral length | 25.2 | 22.2–27.4 | 22.9±1.4 |
| Dorsal-fin base length | 15.1 | 10.7–15.1 | 12.3±1.2 |
| Dorsal-fin spine length | 13.9 | 12.6–16.2 | 13.6±1.1 |
| Anal-fin base length | 14.3 | 12.1–15.9 | 13.1±1.4 |
| Pelvic-fin length | 16.2 | 13.5–16.6 | 14.4±1.1 |
| Pectoral-fin length | 24.5 | 18.3–24.5 | 20.3±1.6 |
| Pectoral-fin spine length | 17.5 | 13.9–19.4 | 15.6±1.8 |
| Caudal-fin length | 24.8 | 21.9–28.8 | 24.2±2.2 |
| Adipose-fin base Length | 13.9 | 12.0–16.5 | 13.1±1.5 |
| Dorsal to adipose distance | 20.4 | 17.1–23.6 | 18.8±1.7 |
| Post-adipose distance | 17.2 | 15.5–18.4 | 15.7±0.9 |
| Caudal peduncle length | 24.3 | 16.1–27.7 | 18.0±3.3 |
| Caudal peduncle depth | 8.9 | 7.8–9.9 | 8.3±0.7 |
| Body depth at anus | 15.4 | 12.4–16.8 | 13.7±1.1 |
| Head length | 30.7 | 26.4–30.7 | 27.0±1.4 |
| Head width at pectoral fin origin | 21.4 | 20.1–23.2 | 20.6±0.8 |
| Head depth just posterior to eye | 14.8 | 9.9–15.2 | 12.7±1.5 |
| %HL |  |  |  |
| Snout length | 48.8 | 34–58 | 46.2±6.1 |
| Interorbital distance | 31.6 | 27–39 | 30.6±3.2 |
| Eye diameter | 10.1 | 10–15 | 10.7±1.3 |
| Nasal barbel length | 16.3 | 10–19 | 13.8±2.3 |
| Maxillary barbel length | 52.1 | 46–66 | 55.3±5.4 |
| Inner mandibular barbel length | 24.7 | 20–36 | 26.4±4.3 |
| Outer mandibular barbel length | 34.8 | 35–54 | 41.0±5.2 |

**Table 2.** Meristic data of *Pseudolaguvia moinensis* (n=13)

| Characters | Holotype | Paratypes |
| --- | --- | --- |
| Dorsal-fin rays | I6 | I6(5), I6½(5), I5½(2) |
| Pectoral-fin rays | I7 | I7(7), I6(1), I7½(4) |
| Pelvic-fin rays | i5 | i5(12) |
| Anal-fin rays | V3½ | iii6(2), iii6½(1), iii7(2), iii5, iii5½,<br>iv5½(3), v4, v4½ |
| Caudal-fin rays | i77i | i77i (12) |
| Pectoral-spine (anterior serrae) | 9 | 9-14 |
| Pectoral-spine (posterior serrae) | 7 | 6(9), 7(3) |
| Procurrent rays (Upper) | 11 | 11(7), 12(2), 8(2), 9(1) |
| Procurrent rays (Lower) | 9 | 9(4), 8(4), 6(2), 11(2) |

Head large and dorsoventrally depressed, degree of depression increasing towards snout margin, lateral profile gently sloping towards snout tip, side and dorsum scattered with numerous small keratinized tubercles, those on occipital region minute. Snout margin moderately rounded. Humero-cubital and scapular processes well-developed. Eye small, ovoid, located entirely on dorsal half of head, situated slightly closer to end of operculum than snout tip. Body also with tubercles but somewhat indistinct. Rayed portions of fins without tubercles.

Mouth subterminal, consisting of fleshy papillated lips, region between lower lip and thoracic adhesive apparatus densely papillated, upper jaw projecting over lower jaw, premaxillary tooth band partially visible when mouth closed. Upper lip continuing into maxillary barbel, and barbel bases connected to sides of head through small triangular dermal flap at base, just above corner of mouth.

Gill opening wide, extending obliquely from posttemporal to isthmus. Supraoccipital spine terminating very close to nuchal shield. Weberian lamina well developed and situated either side of supraoccipital spine, extending parallel or sometime slightly radiating away. Lateral line complete and midlateral with about 65–75 pores. Total vertebrae 30–31 in four specimens. Caudal peduncle short and moderately slender. Anus situated mid of pelvic and anal fin. Skin prominently tuberculate on dorso-lateral side of head, moderately tuberculate on dorsal third of body. Abdomen with thoracic adhesive apparatus, anteriorly broader, gently reduced to narrow posteriorly, consisting of longitudinal, unculiferous ridges arranged in an elliptical field, and with prominent central median depression, about twice as long as broad, extending close to pelvic-fin base (Fig. 3), posterior most ridges small and weakly developed. Nostril situated in between eye and tip of snout, anterior nare separated by short nasal barbel, dividing it into two parts; anterior nare more smoothly rounded than posterior.

**FIGURE 3.**
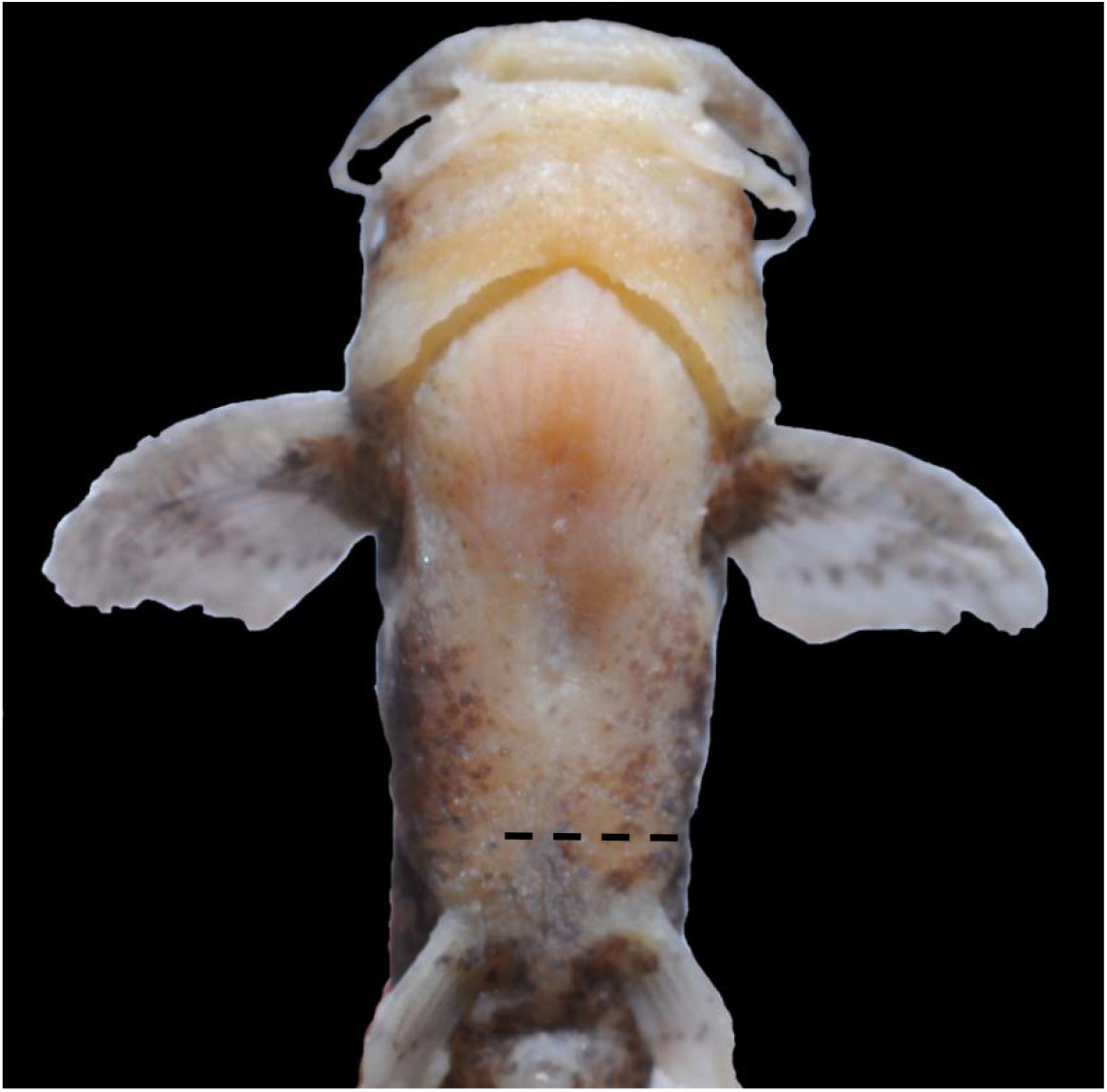
*Pseudolaguvia moinensis*, RGUMF 1897, holotype, showing extension of adhesive apparatus closer to pelvic fin bases.

Barbels in four pairs (one pair each of nasal, maxillary, outer and inner mandibular), nasal barbel very short with broad base and pointed tip, arising from the internarial septum, reaching only two thirds the distance from naris to eye. Maxillary barbel slender, with broad skin flap at base, dorsoventrally depressed and mostly extending onto posterior orbital margin or slightly exceeding, but not reaching pectoral-spine origin. Outer mandibular barbel with thick base, moderately depressed and extending closer to pectoral-spine origin; inner mandibular barbel shorter and cylindrical, reaching or closer to isthmus.

Dorsal fin located about two-fifths along body, with 1 spine and 5½ (2), 6*(6), or 6½ (5) branched rays. Dorsal-fin spine short, robust, straight, moderately compressed; adpressed spine longitudinally extending to anterior margin of creamy band or slightly beyond pelvic-fin origin; anterior margin of spine smooth, posterior margin with 3–4 indistinct serrations. Adipose fin short, roughly triangular, its anterior margin straight and slightly longer than posterior, posterior margin mostly straight or rarely slightly convex with incised posterior end, originating slightly anterior or vertical to anal-fin origin.

Pectoral fin with sharp pointed spine, extending very close to pelvic-fin origin, anterior and posterior margin gently arched and with 1 spine and 7*(8), 6(1), 7½(4) branched rays; anterior spine margin with 9*(3), 11(5), 12(3) or 13(2) small serrations, distally directed, size decreasing towards base, proximal most serrae small and granulated; posterior spine margin with 6(9)–7*(3) distinct serrations, distantly located and proximally directed. Pectoral girdle with prominent postcoracoid processes, hidden beneath skin, extending to midway between base of last pectoral-fin ray and its tip. Pelvic fin soft, with 1 unbranched and 5 (13) branched rays, located almost at middle of body; origin perpendicular to base of last second ray of dorsal fin, anterior margin slightly convex, posterior obliquely straight; first or second branched rays longest; adpressed fin extending beyond anus, but never reaching anal-fin origin; tip of last ray reaching middle to posterior margin of anus in adult specimens, extending slightly past anus in juveniles. Anal fin soft, with diverse rays pattern– iii6 (2), iii5, iii5½, iii6½, iii7 (2), iv5½ (3), v3½*, v4, or v4½ rays, anterior margin slightly convex and distal obliquely straight, adpressed tip reaching posterior margin of oblique band on caudal peduncle. Caudal fin forked, longer than all other fins, and with i77i principal rays; upper and lower lobes subequal, lower slightly deeper than upper and sometime slightly longer, tip of upper lobe more pointed than lower. Procurrent rays symmetrical, extending anteriorly only to hypural margin, upper lobe with 8(2), 9(1)11* (7) or 12(2) rays and lower with 6(2), 8(4), 9*(4) or 11(2).

Other equivalent characters: predorsal length almost equals distance between dorsal-fin origin and mid of adipose fin; body depth at anus equals adipose-fin base length; dorsal-fin spine equals anal-fin base length; post adipose distance equals anal-fin base length.

#### Coloration in preservative

In 75% alcohol: Dorsolateral surface of head and body light brown to dark brown, darker on occipital region. Head and body with minute, brown to darkbrown spots scattered throughout, sparsely scattered on ventral creamy region between snout tip and pelvic-fin origin. Two light brown to cream bands on side of body– one band situated between dorsal and adipose fins and another on caudal peduncle, former band V-shaped, vertex pointing posteriorly, broader than latter, not contiguous dorsally and ventrally, latter band, indistinct and sometime distinct, obliquely positioned straight to posterior margin of adipose fin. A single pair of creamy spots on body just below dorsal-fin origin. Dorsal fin brown with hyaline distal margin. Adipose fin brown, with hyaline anterior and posterior distal margin. Caudal fin with two hyaline medial bands on each lobe, with complete anterior end, touching outer margin of each lobe, and tip region of each lobe also hyaline. Caudal fin base dark brown, extending over median rays, and then bifurcating into two thin stripes that confluence with large dark brown patch on each lobe, overall giving appearance of W-shaped. Pectoral, pelvic fins hyaline except small light brown bases and narrow subdistal streaks. Anal fin with distinct brown base and light brown subdistal streak. Maxillary and outer mandibular barbels creamy ventrally, light brown dorsally, annulated with 3-4 brown rings in the former and 2-3 in the latter, more prominent in live. Nasal barbel dorsally light brown and ventrally cream, and inner mandibular barbels mostly creamy.

#### Distribution and habitat

Presently known only from the Moin drainage, a small tributary draining into Poma River towards west, in Papum Pare District, Arunachal Pradesh, India (Fig. 5). Further, the Poma River flows west and south along the dense forests, over rocks, sand and gravelly substratum and enters the sandy plain of the neighboring state, Assam, which, eventually merge with Brahmaputra River in the south. The water was moderately flowing and clear/muddy in two different occasions, and bed consisting of small to medium-sized stones, pebbles, cobbles, large boulders, and sand (Fig. 4). It has been noted that fish camouflages with habitat pattern and water condition— Dorsal and lateral surfaces of body and vertical bands along flank changes to light brown in muddy water, whereas brown to dark brown with cream bands in clear water with various colored substrates. The riparian vegetation comprised of shrubs and small to large trees towards uphill slope. No other species of *Pseudolaguvia* were encountered syntopically. Other associated fishes in the collection site or nearby were: (Cyprindiae) *Opsarius bendelisis*, *Opsarius barna*, *Devario aequipinnatus*, *Devario devario*, *Danio dangila*, *Garra annandalei*, *Garra birostris*, *Tor putitora*; (Nemacheilidae) *Aborichthys uniobarensis*; (Psilorhynchidae) *Psilorhynchus balitora*; (Badidae) *Badis badis*; (Channidae) *Channa pomanensis*; (Amblycepitidae) *Amblyceps arunachalensis*, *Amblyceps apangi*; and (Olyridae) *Olyra longicaudata*.

**FIGURE 4.**
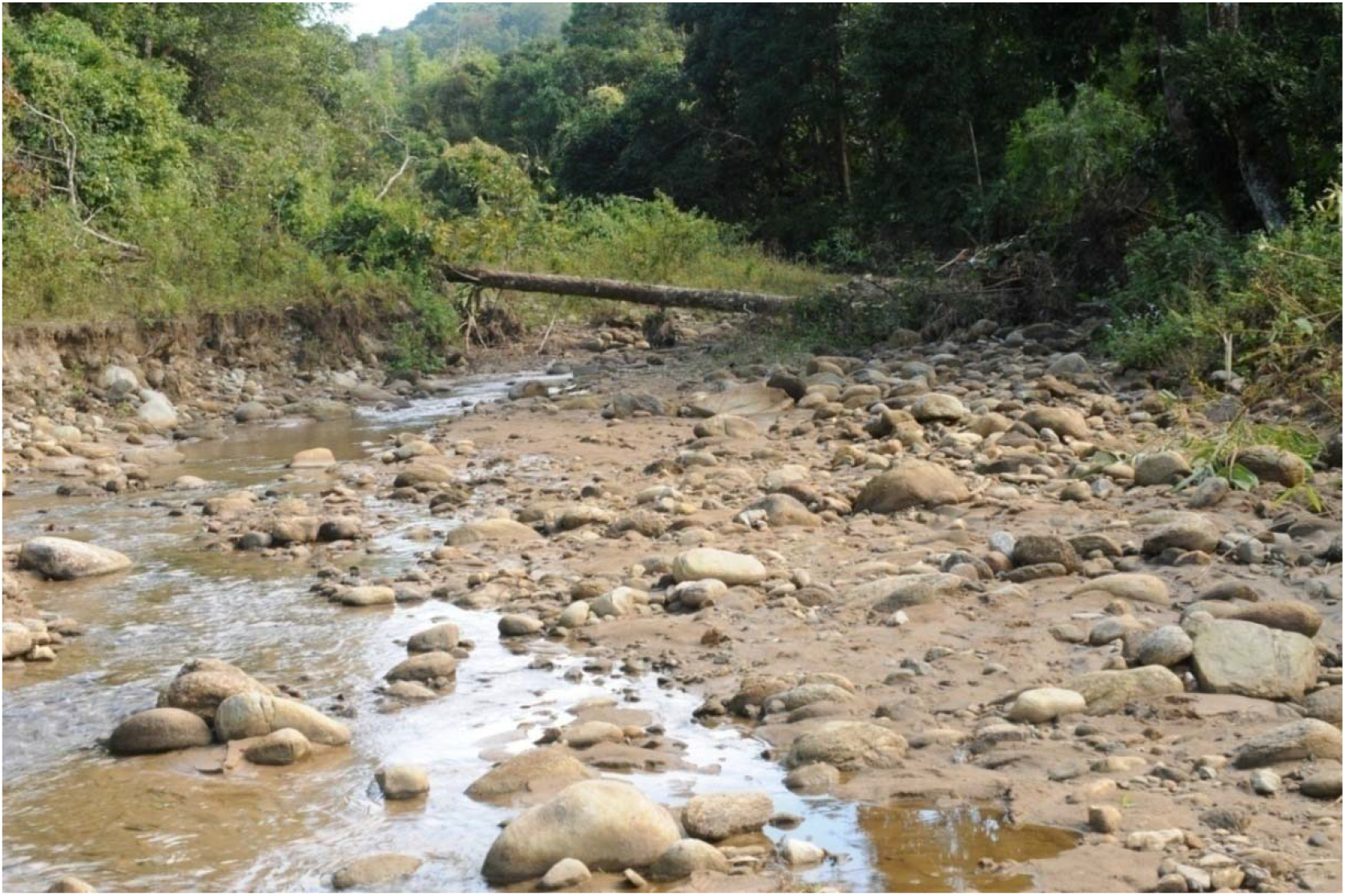
Habitat of *Pseudolaguvia moinensis*, Moin drainage, a tributary of Poma River (Brahmaputra River basin), Arunachal Pradesh, India.

**FIGURE 5.**
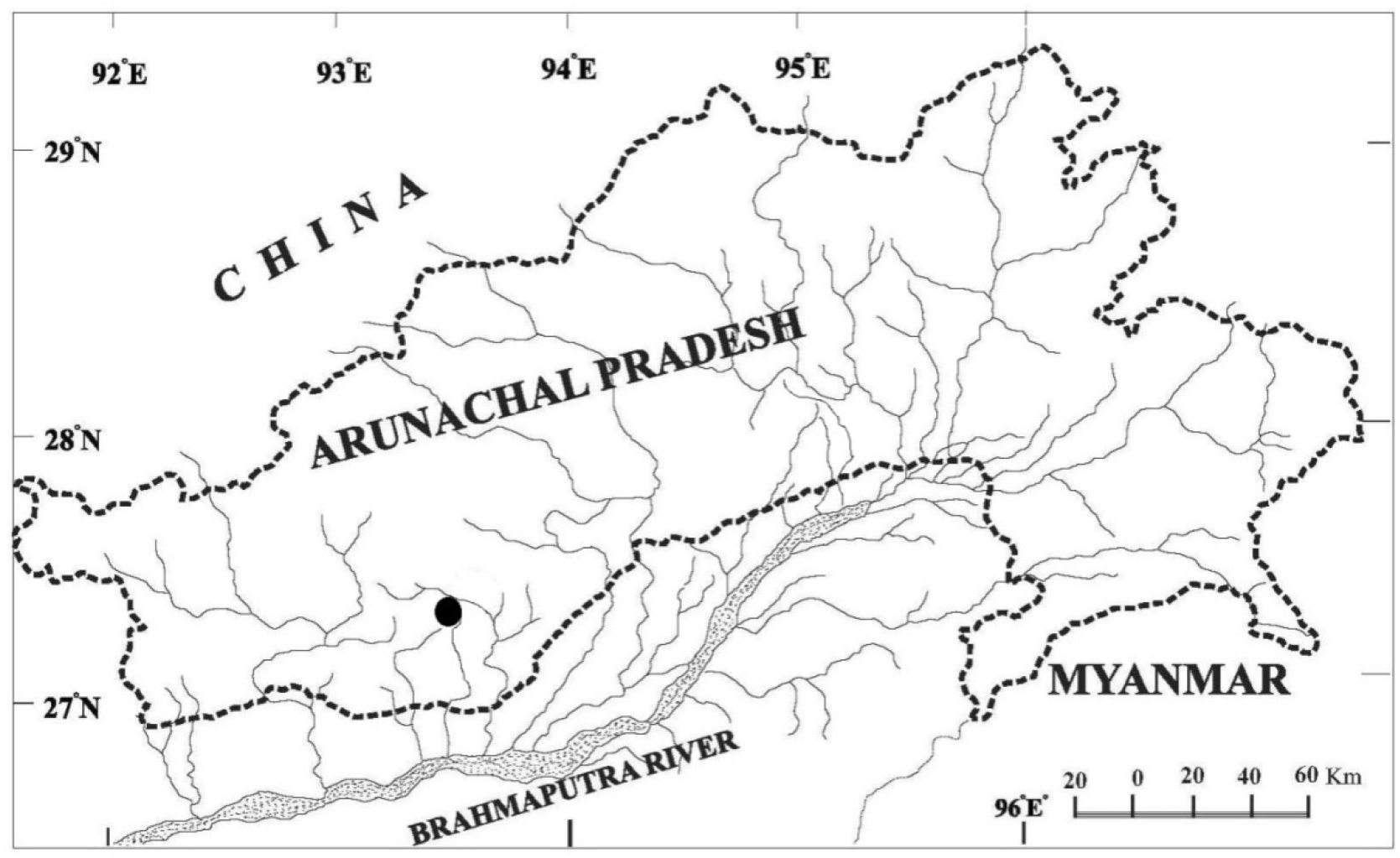
Drainage map of Arunachal Pradesh, filled circle showing type locality of *Pseudolaguvia moinensis*; near Moin village, Papum pare District.

#### Etymology

*Pseudolaguvia moinensis* is named for the ‘Moin drainage’ a small tributary of Poma River, an adjective.

## Discussion

Of the total twenty three recognized species of *Pseudolaguvia*, about half (9 species) are confined to Brahmaputra River drainages in northeastern India and Rangapani Khal, Bangladesh (Ng, 2009; Ng & Tamang, 2012; Tamang & Sinha, 2014): *P. foveolata*, *P. ferula*, *P. viriosa*, *P. muricata*, *P. magna*, *P. jiyaensis*, *P. ribeiroi*, *P. shawi*, and *P. ferruginea*; seven species *P. kapuri*, *P. inornata*, *P. flavida*, *P. virgulata*, *P. assula*, *P. nepalensis* and *P. flavipinna* from the Ganga drainage system (Tamang & Sinha, 2014; Rayamajhi *et al*., 2016; Bhakat, 2019); two species each: *P. nubila* from the Kaladan River, and *P. fucosa* from the Karnaphuli River drainage, Mizoram (Ng *et al*., 2016); *P. austrina* from Bharathapuzha River and *P. lapillicola* from Subramanya and Kumaradhara River in peninsular India (Britz *et al*., 2013); and *P. tenebricosa* and *P. tuberculata* from the Sittang and Irrawaddy river systems, Myanmar (Britz, 2003); and lastly, one species *P. spicula* from the Surma-Meghna river drainage in northeast India and Bangladesh (Ng & Lalramliana, 2010b).

The new species is distinguished from all congeners except *P. ferruginea* and *P. focusa* by having an ovoid to elongate light brown to cream marks either side on ventro-lateral margin just above anus, and except for *P. ribeiroi*, *P. ferruginea*, *P. foveolata*, *P. muricata*, *P. kapuri, P*. *jiyaensis*, *P. flavipinna*, *P. nepalensis*, *P. nubila*, *P. spicula* and *P. tuberculata* in having a W-shaped dark brown band on caudal fin (vs. absent). Further, *Pseudolaguvia moinensis* can be differentiated from the seven species *P. ribeiroi*, *P. fucosa*, *P. flavida*, *P. kapuri*, *P. muricata*, *P. nepalensis*, and *P. virgulata*, which commonly shares the presence of serrations on the anterior edge of the dorsal spine (Ng *et al*., 2016), whereas smooth in case of *P. moinensis*.

Congeners from Brahmaputra River system: *Pseudolaguvia moinensis* is distinguished from *P. foveolata* in having a longer thoracic adhesive apparatus (reaching beyond vs. not reaching base of last pectoral-fin ray), shorter pelvic fin (13.5–16.6% SL vs. 19.0), adipose-fin base (12.0–16.5% SL vs. 24.0), and deeper caudal peduncle (7.8–9.9% SL vs. 5.0); from *P. ferula*, *P. viriosa* and *P. muricata* by a shorter dorsal-fin spine (12.6–16.2% SL vs. 17.3–18.7 in *P. ferula*; 23.4–29.0 in *P. viriosa*; 21.2–26.7 in *P. muricata)*, and except for *P. ferula* by a shorter pectoral-fin spine (13.9–19.4% SL vs. 26.9–32.9 in *P. viriosa* and 26.8–35.7 in *P. muricata*); from *P. magna* by having (vs. lacking) light brown to cream band on the body posterior to dorsal fin, lacking (vs. having) a roughly rectangular to ovoid pale to cream patch on the mid-dorsal region of the body, a longer adhesive apparatus (extending vs. not extending) beyond last pectoral fin ray; from *P. jiyaensis*, *P. ribeiroi*, and *P. shawi* by having a V-shaped (vs. vertical) band on side of the body, bands on either side not contiguous (vs. contiguous) dorsally; further: from *P. jiyaensis* by having a shorter adhesive apparatus (not reaching vs. reaching) pelvic-fin base; and from *P. ribeiroi* by a deeper caudal peduncle (7.8–9.9% SL vs. 6.6–7.2).

Among all congeners, *Pseudolaguvia moinensis* shares *P. ferruginea*, the light brown to cream V-shaped and an oblique bands along the flank of the body, but differs by a deeper body at anus (12.4–16.8% SL vs.10.5–12.4), anal fin with a narrower (maximal width 5–8½ times anal-fin length vs. slightly more than 2½; data calculated from Ng, 2009; fig.7b, holotype) subdistal brown band; and tip of anal fin extending beyond (vs. extending at the level of) posterior end of adipose fin.

Congeners from Ganges River system: *Pseudolaguvia moinensis* is distinguished from congeners *P. inornata*, *P. virgulata* and *P. assula* by the presence (vs. absence) of light brown to cream band on side of the body; further: from *P. inornata* by having a shorter maxillary (46–65% HL vs. 78–83), a shorter dorsal- (12.6–16.2% SL vs. 18.6–21.7) and pectoral-fin spines (13.9–19.4% SL vs. 20.4–23.3); from *P. inornata* and *P. virgulata* by a shallower head (depth 9.9–15.2% SL vs. 15.9–19.4); from *P. assula* by a shorter pectoral- (13.9–19.4% SL vs. 23.3–28.3) and dorsal-fin spines (12.6–16.2% SL vs. 20.3–24.8); from *P. flavida* by having a longer pelvic fin (13.5–16.6% SL vs. 10.2), prepectoral (22.2–27.4% SL vs. 19.5), and a deeper body at anus (12.4–16.8% SL vs. 11); from *P. kapuri* by a shorter adipose-fn (12.0-16.5% SL vs. 17.1–18.9), V-shaped (vs. almost vertical) band on the side of the body, which is not contiguous (vs. contiguous) dorsally, and caudal fin median rays dark brown (vs. hyaline); from *P. nepalensis* in having a shorter head (26.4–30.7% SL vs. 32.4–33.6), prepectoral (22.2–27.4% SL vs. 28.4–29.9), dorsal-fin spine (12.6–16.2% SL vs. 22.7–24.4), pectoral fin (18.3–24.5% SL vs. 28.9-35.2), and a longer adhesive apparatus (extending vs. not extending) beyond last pectoral-fin ray; and from *P. flavipinna* by lacking (vs. having) V-shaped yellowish hyaline band on caudal fin, in having a shorter predorsal (27.0–31.7% SL vs. 44.0-46.4), prepelvic (47.1–52.8% SL vs. 54.5-56.4), shorter caudal fin (21.9–28.8% SL vs. 31.2-32.4), and longer and deeper caudal peduncle (length 16.1–27.7% SL vs. 13.0-14.5 and depth 7.8–9.9% SL vs. 12.6-14.4).

Species from Kaladan and Karnaphuli River drainages: *Pseudolaguvia moinensis* is distinguished from *P. nubila* by having a shorter dorsal-fin base (10.7–15.1% SL vs. 15.1–17.3), shorter dorsal spine (12.6–16.2% SL vs. 16.4–19.3), and a shallower head (depth 9.9–15.2% SL vs. 15.3–18.9); and from *P. fucosa* by a shorter dorsal- (12.6–16.2% SL vs. 18.0–21.6) and pectoral-fin spines (13.9–19.4% SL vs. 20.7–26.1), a shorter pectoral fin (18.3–24.5% SL vs. 24.6–29.5), adipose-fin base (12.0–16.5% SL vs. 21.5–26.3) and absence (vs. presence) of a pale, y-shaped marking on the dorsal surface of the head, and caudal fin with (vs. without) hyaline median cross band.

*Pseudolaguvia moinensis* is distinguished from *P. spicula*, a congener from the Surma-Meghna River system by having an anterior end of medial hyaline bands on each lobe of caudal fin complete (vs. incomplete), reaching (vs. not reaching) outer margin of each lobe.

Until now, only two species each are known from peninsular India and Myanmar: *P. austrina* from the Bharathapuzha River and *P. lapillicola* from the Subramanya and Kumaradhara Rivers, while *P. tenebricosa* and *P. tuberculata* from the Sittang and Irrawaddy river systems. *Pseudolaguvia moinensis* is distinguished from *P. austrina* and *P. lapillicola* by a narrower head (20.1–23.2% SL vs. 24.2–27.0 in *P. austrina*; 24.6–25.9 in *P. lapillicola)*, shorter nasal barbel (10–19% HL vs. 25–47), and fewer vertebrae (30–31 vs. 33–34); further: from *P. austrina* by having fewer (6-7 vs. 9-12) serrae on posterior margin of pectoral spine, a longer and shallower caudal peduncle (length 16.1–27.7% SL vs. 8.8–14.7 and depth 7.8–9.9% SL vs. 10.3– 14.7); from *P. lapillicola* by a shallower head (depth 9.9–15.2% SL vs. 19.4–20.7), and a shorter caudal fin (21.9–28.8% SL vs. 28.6–30.3). It can be further distinguished externally by having thoracic adhesive apparatus extending beyond (vs. middle) of last pectoral fin ray, and without (vs. with) prominent dark brown to black spots on body; from *P. tenebricosa* by having a shorter pelvic fin (13.5–16.6% SL vs. 16.0–20.9), and caudal fin (20.6–25.9% SL vs. 25.7–43.9), a shallower caudal peduncle (7.8–9.9% SL vs. 10.3–14.7), and anterior end of medial hyaline bands on each lobe of caudal fin complete (vs. incomplete); and from *P. tuberculata* by having the adipose fin not contiguous (vs. contiguous) with the dorsal-fin base.

## Comparative material

### Pseudolaguvia viriosa

ZSI V/APRC/P-524 (holotype), 26.0 mm SL; RGUMF 007 (13 paratypes), 23.1–27.2 mm SL India: Arunachal Pradesh, East Siang district, Sille River, approximately 1 km upstream from RCC bridge, about 10 km from Ruksin and about 26 km before Pasighat, 27°52’37.6”N 95°18’18.0”E.

### P. tuberculata

ZSI F10876/1 (holotype), 29.6 mm SL; Myanmar: Myitkyina, Sankha, a large hill-stream, midway between Kamaing and Mogaung. Additional data from Britz& Ferraris (2003).

### P. shawi

ZSI F 10085/1 (holotype); India: West Bengal, Mahanadi River below Darjeeling [Brahmaputra drainage]. Additional data from Ng (2009) and Ng (2005a).

### P. magna

ZSI/ V/APRC/P-947 (holotype); 46.1 mm SL; ZSI/ V/APRC/P-948 (8 paratypes), 31.9–46.7 mm SL, (2) 40.5–42.3 mm SL (skeleton), India: Arunachal Pradesh, Lower Dibang Valley District, Jiya stream, near Bolik village, approximately 14 km from Roing towards Shantipur, Assam, 28°00.377’N 95°45.562’E; 149 m asl.

### P. jiyaensis

ZSI/V/APRC/P-1034 (holotype); 29.9 mm SL; ZSI/V/APRC/P-1035 (15 paratypes) 25.6–31.2 mm SL, India: Arunachal Pradesh, Lower Dibang Valley District, Jiya stream, near Bolik village, approximately 14 km from Roing towards Shantipur, Assam, 28°00.377’N 95°45.562’E; 149 m asl.

#### Other source of comparison

*Pseudolaguvia foveolata*, *P. kapuri*, *P. ribeiroi*, *P. ferruginea* and *P. flavida*: Ng (2005a) and Ng (2009). *P. ferula*: Ng (2006a), *P. inornata* and *P. muricata*: Ng (2005b), *P. tenebricosa, P. tuberculata*: Britz & Ferraris (2003), *P. virgulata*: Ng & Lalramliana (2010a), *P. spicula*: Ng & Lalramliana (2010b). *P. austrina*: Radhakrishnan *et al*. (2011), *P. lapillicola*: Britz *et al*. (2013), *P. assula*: Ng & Conway (2013), *P. nubila*: Ng *et al*., 2013, *P. flavipinna*: Bhakat, 2019, *P. focusa*: Ng *et al*., 2016, *P. nepalensis*: Rayamajhi *et al*., 2016.

## Acknowledgments

We are grateful to the Hon’ble, Vice Chancellor, Rajiv Gandhi University, Rono Hills, Doimukh for providing infrastructure facilities. We also thank A.K. Karmakar (ZSI, Kolkata) for access to material (holotype of *P. tuberculata* and *P. shawi)* in their care for comparative study. A thanks also goes to Mihin Amo, Krima Queen Machahary and Tok Loma for their assistance in the field. This work was funded by support from the DBT New Delhi sponsored project (BT/PR116506/NER/95/210/2015) and is gratefully acknowledged.

